# ASFV pD345L protein negatively regulates NF-κB signaling through inhibiting IKK kinase activity

**DOI:** 10.1101/2021.09.06.459096

**Authors:** Huan Chen, Zhenzhong Wang, Xiaoyu Gao, Jiaxuan Lv, Yongxin Hu, Yong-Sam Jung, Shanyuan Zhu, Xiaodong Wu, Yingjuan Qian, Jianjun Dai

**Author notes:** Corresponding Author: Yingjuan Qian, or Xiaodong Wu.

## Abstract

NF-κB is a critical transcription factor in immediate early viral infection, including African swine fever virus (ASFV), playing an important role in inflammation response and expression of antiviral genes. ASFV encodes for more than 151 proteins by its own transcription machinery and possesses a great capacity to evade or subvert antiviral innate immune responses. A couple of such viral proteins have been reported, but many remain unknown. Here, we showed that pD345L, an ASFV-encoded lambda-like exonuclease, is an inhibitor of cGAS/STING mediated NF-κB signaling by blocking IKKα/β kinase activity. Specifically, we showed that overexpression of pD345L suppresses cGAS/STING induced IFNβ and NF-κB activation, resulting in decreased transcription of IFNβ and several pro-inflammatory cytokines, including IL-1α, IL-6, IL-8, and TNFα. In addition, we showed that pD345L targeted at or downstream of IKK and upstream of p65. Importantly, we found that pD345L associates with KD and HLH domains of IKKα and LZ domain of IKKβ, and thus interrupts their kinase activity on downstream substrate IκBα. Finally, we showed that pD345L inhibition of NF-κB signaling was independent of its exonuclease activity. Taken together, we concluded that pD345L blocks IKK α/β kinase activity by protein-protein interaction and thus disrupts cGAS/STING mediated NF-κB signaling.

**Importance:** African Swine Fever (ASF) is one of the most devastating and economically significant swine diseases caused by African swine fever virus (ASFV). Since expanding of ASFV affected areas into Asian countries, especially China, an effective vaccine is urgently needed more than ever. Therefore, it is of great significance to understand the interaction between ASFV infection and host immune responses. The NF-κB signaling plays a central role in innate and acquired immune responses. Activation of IκB kinase (IKK) complex is a key step of both canonical and non-canonical NF-κB pathways, and is commonly targeted by different viruses. But no ASFV protein has been shown to regulate IKK yet. In this study, we demonstrated that pD345L blocks IKKα/β kinase activity by protein-protein interaction and thus disrupts cGAS/STING mediated NF-κB signaling. It has been shown that conventional vaccine development approaches are proven to be inapplicable to ASFV. Neither subunit nor DNA vaccine provides efficient protection. Gene deleted live-attenuated vaccine candidates render adequate protection, but establishment of chronic or persistent infection in vaccinated animals and risk of recombination with filed strains are key challenges. To overcome these, one potential strategy would be generation of replication-defective viruses. As a lambda-like exonuclease, pD345L plays a critical role in ASFV replication and ASFV deficient in D345L cannot be rescued. Given the dual role of pD345L in virus replication and immune evasion, it may serve as a potential target for replication-defective virus vaccine development.

## Introduction

African Swine Fever (ASF) is one of the most devastating and economically significant swine diseases caused by African swine fever virus (ASFV), a large enveloped double-stranded DNA virus that infects domestic pigs (*Sus scrofa domestica*) and wild boars (*Sus scrofa*) with a morbidity and mortality rate up to 100%. Since the first report in Kenya in 1921, ASFV genotype I and genotype II have escaped from Africa to Europe, South America, the Caucasus, and Russian Federation (1). ASFV genotype I was successfully eradiated in all countries outside Africa except Italian sardinia island. In August 2018, ASFV genotype II was introduced into China, the largest pork producer and consumer in the world, and rapidly transmitted to almost all Chinese provinces as well as more than ten Asian countries, including Mongolia, Korea, Vietnam, Laos, Cambodia, Myanmar, the Philippines, Indonesia, Timor-Leste and Papua New Guinea (2). As a result, millions of pigs were culled and pig inventory in these countries was significantly decreased. It not only largely threatened the swine industry and heparin supply worldwide but also caused tremendous social and economic impacts(3). However, there are no effective vaccines or antivirals available yet. Therefore, it’s of great importance to understand ASFV and host interaction to find new clues for vaccine development.

ASFV, the sole member of the family Asfarviridae and the only known DNA arbovirus, which shares a common origin with nucleocytoplasmic large DNA virus (NCLDVs), including poxviruses, iridoviruses, and mimiviruses (4, 5), accomplishes its replication cycle and assembles newly synthesized virions in the cytoplasmic viral factory. ASFV virion is an icosahedral particle with a double-stranded DNA genome of 170-190 kbp that contains 151 to 167 open reading frames (ORFs) depending on viral strains (6). Except structural proteins, it also encodes a variety of proteins involved in gene transcription, replication, nucleotide metabolism, DNA repair and immune regulation (7). ASFV primarily targets host monocyte-macrophage lineage cells, but only replicate with a very low efficiency and easy to loss of immunogenicity in immortalized cell lines. As innate immune cells, macrophages have a comparably harsh intracellular environment for engulfed microorganisms. To complete its replication cycle, ASFV should possess a delicate mechanism to modulate and thus generate a favorable intracellular condition. For example, macrophages produce reactive oxygen species (ROS) in response to virus infection which not only creates oxidative DNA lesions destroying the viral genome integrity (8, 9), but also inhibits viral capsid assembly and maturation (10). Base excision repair (BER) pathway is the primary repair system in response to ROS-induced DNA damage. Interestingly, except DNA glycosylase, ASFV encodes for all BER pathway enzymes, including AP endonuclease (pE296R), PCNA-like (pE301R), DNA polymerase X-like (pO174L), Lambda-like exonuclease (pD345L) and DNA ligase (pNP419L). It was reported that pE296R is required for ASFV growth in macrophages but not in Vero cells (11), and pD345L is indispensable for ASFV propagation (12), suggesting an essential role of DNA repair system in ASFV replication.

In addition, as a large DNA virus, ASFV possesses multiple strategies to evade host innate immune defenses usually initiated by recognition of pathogen-associated molecular patterns (PAMPs) by specific pattern recognition receptors (PRRs), such as cyclic GMP-AMP synthase (cGAS), a recent identified cytosolic DNA sensor. cGAS recognizes dsDNA and synthesizes the second message cGAMP which associates with and activates the endoplasmic reticulum (ER)-localized adaptor molecule Stimulator of Interferon Genes (STING). Once activated, STING translocates from ER to Golgi where it phosphorylates TBK1 and IKK to initiate either IRF3- or NF-κB-mediated innate immune responses. The NF-κB signaling plays a critical role in regulating immune and inflammatory responses through canonical and non-canonical pathways (13). The canonical pathway is activated by multiple stimuli, such as host-derived cytokines, viral or bacterial products and stress-related factors, leading to IKK complex mediated phosphorylation and degradation of IκBα and thus releasing p65/p50 dimers. IKK complex contains two catalytic subunits, IKKα and IKKβ, and one regulatory subunit IKKγ (NEMO). Study also showed a new role of IKKα in phosphorylating histone H3, which is critical for the activation of NF-κB-controlled gene expression (14, 15). The non-canonical pathway is activated by a specific subset of TNF receptors, including CD40, lymphotoxin β receptor and RANK, which requires NIK and IKKα mediated p100 processing to p52 (16). Generally, IKK acts as a key regulator in NF-κB-mediated anti-viral response, and thus IKK is an important target for virus immune evasion.

A few ASFV proteins have been identified to regulate host immune responses. For examples, A238L, an IκB homologue, is able to either prevent NF-κB binding with its responsive elements of the target gene promoters (17) or suppress the transcriptional activity of NF-κB via interacting with p300/CBP (18). Both I329L, a viral TLR homologue, and A528R inhibit IRF3 and NF-κB activation (19). In addition, DP96R was discovered to inhibit the cGAS-STING signaling pathway via targeting TBK1 and IKKβ with an unknown mechanism (20). Recently, MGF-505-7R has been shown to suppress the cGAS-STING pathway via promoting autophagy-mediated STING degradation (21). Moreover, multigene family 360/530 (MGF360/530) deletion virus shows attenuated virulence and enhanced type I IFN production (22, 23). Chronic/persist infection caused by live attenuated viruses indicates that additional ASFV proteins modulating host immune response remain to be identified.

Importantly, here we found that pD345L plays a negative role in cGAS/STING-mediated NF-κB signaling pathway independent of its exonuclease activity, but altering IKKα and IKKβ kinase activity through protein-protein interaction. These suggest that pD345L is important for virus replication in host cells through its DNA repair and immune suppression activities.

## Results

### ASFV pD345L attenuates IFNβ production through cGAS/STING mediated NF-κB signaling

Upon ASFV infection, cytosolic DNA will be mainly detected by the key DNA sensor cGAS (24), then activating STING-dependent type I interferon response. To identify ASFV proteins that regulate cGAS/STING-mediated immune response, a number of ASFV-encoded proteins were screened using dual luciferase reporter assay by transfection of each viral protein with the INFβ, NF-κB or IRF3 luciferase reporter, designated INFβ-Luc, NF-κB-Luc, or IRF3-Luc, along with co-transfection or treatment with innate immune stimulators. Among these, pD345L was identified. To confirm this, a dual-luciferase reporter assay was conducted with co-transfection of the IFNβ-Luc along with cGAS, STING and/or D345L expression vectors, and results showed that pD345L obviously decreased cGAS/STING-induced activation of the IFNβ promoter in a dose-dependent manner (Figure 1A and 1B). However, pD345L did not change poly(I:C)-stimulated activation of the IFNβ promoter (Figure 1C). Next, to further determine whether NF-κB or IRF3 pathway is targeted by pD345L, the effect of pD345L on the NF-κB and IRF3 promoter activation was analyzed. Results showed that pD345L suppressed cGAS/STING-induced NF-κB-Luc activity (Figure 1D), but not IRF3-Luc activity (Figure 1E). Similarly, pD345L also inhibited NF-κB-Luc induced by TNFα, which activates the IKK/NF-κB signaling (Figure 1F). These findings suggest that pD345L selectively inhibits cGAS/STING-mediated NF-κB signaling pathway.

**FIG 1.**
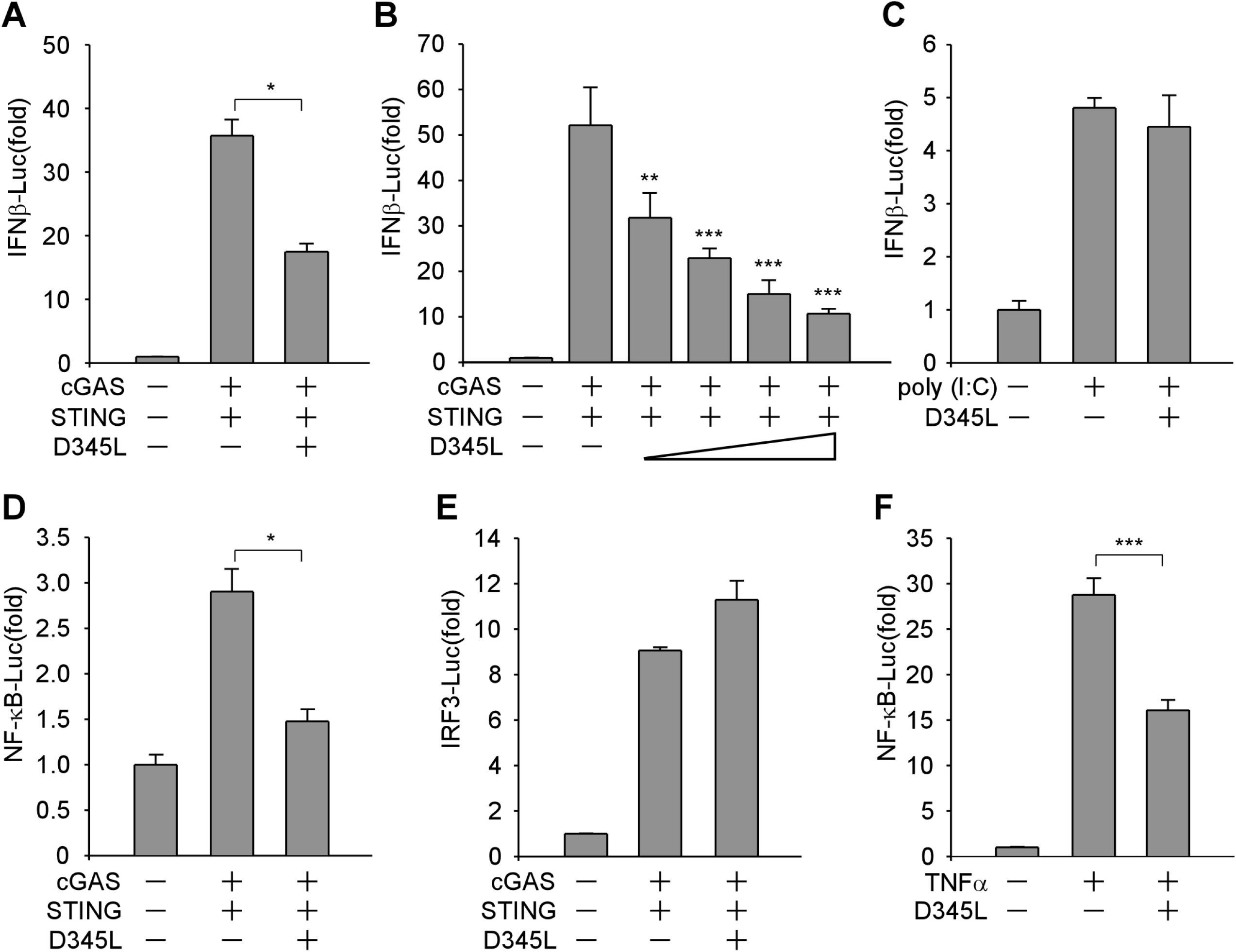
pD345L inhibits IFNβ expression through cGAS-STING mediated NF-κB signaling. (A) 293T cells transfected with pGL3-Basic-IFNβ-Luc and pCMV-RL along with empty vector, pcDNA3.1-HA-cGAS, pcDNA3-2xFlag-STING and/or p3XFLAG-CMV-D345L for 24 hours were collected for measuring luciferase activities. (B) The experiment was performed as panel A except using an increasing amount of p3XFLAG-CMV-D345L. (C) The experiment was performed with PK15 cells transfected with pGL3-Basic-IFNβ-Luc, pCMV-RL and p3XFLAG-CMV-D345L or empty vector for 24 hours followed by treatment with poly (I:C) (10μg/ml) for 12 hours. (D) The experiment was performed as panel A except using NF-κB-Luc. (E) The experiment was performed as panel A except using IRF3-Luc. (F) The experiment was performed with 293T cells transfected with pGL3-Basic-IFNβ-Luc, pCMV-RL and p3XFLAG-CMV-D345L or empty vector for 24 hours followed by treatment with TNFα (40ng/ml) for 6 hours.

### pD345L suppresses the production of IFNβ and pro-inflammatory cytokines

To confirm whether pD345L inhibits IFNβ gene transcription, Real-Time PCR was performed to examine the effect of pD345L on IFNβ mRNA expression levels upon co-transfection of cGAS/STING along with or without D345L in 293T cells. Results showed that upregulation of IFNβ mRNA induced by cGAS/STING was largely reduced in the presence of pD345L (Figure 2A). As a key transcription factor governing the pro-inflammatory signaling pathway(25), NF-κB controls the expression of a set of pro-inflammatory cytokines, including I-L1α, IL-6, IL-8 and TNFα. Therefore, if pD345L regulates NF-κB activity, it will be able to modulate the transcription of other NF-κB downstream targets in addition to IFNβ. To test this, the mRNA levels of IFNβ as well as four pro-inflammatory cytokines were examined in WSL, an immortalized wild boar lung fibroblast cell line, and PK15, a pig kidney cell line, that untreated or treated with 2’3’-cGAMP. Results showed that pD345L decreased not only the mRNA levels of IFNβ in both cells with or without 2’3’-cGAMP stimulation, but also those of IL-1α, IL-6, IL-8 and TNFα (Figure 2B and C). Collectively, these results indicate that pD345L inhibits the cGAS/STING-mediated NF-κB signaling.

**FIG 2.**
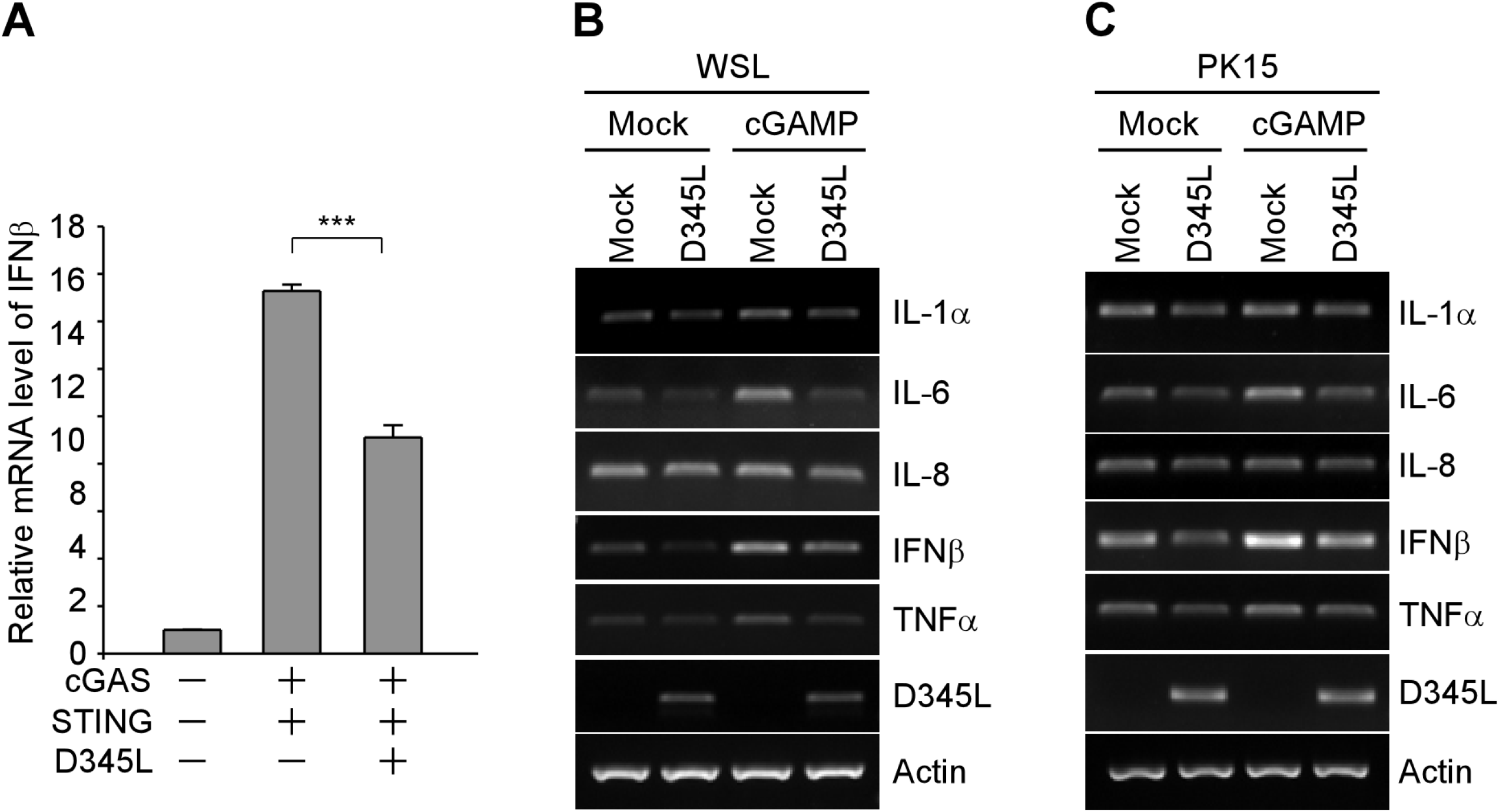
pD345L suppresses mRNA expression levels of IFNβ and pro-inflammatory cytokines. (A) Quantitative real-time PCR was performed with RNA extracted from 293T cells transfected with pcDNA3.1-HA-cGAS and pcDNA3-2xFlag-STING along with p3XFLAG-CMV-D345L or empty vector for 24 hours. (B)WSL or (C) PK15 cells were transfected with p3XFLAG-CMV-D345L or empty vector for 30 hours followed by treatment with 2’3’ cGAMP (5μg/ml) for 4 hours and then harvested for RNA extraction. Semiquantitative RT-PCR assay was carried out to detect IL-1α, IL-6, IL-8, IFNβ, TNFα, D345L and Actin mRNA expression levels.

### pD345L suppresses NF-κB activation at or downstream of IKK and upstream of p65

Upon association of a ligand to its cell surface receptor or recognition of cytosolic DNA by cGAS/STING, adaptors, such as TRAFs, TBK1 or TAK1, will be recruited to the IKK complex to mediate IκBα phosphorylation and degradation, and consequently enable translocation of the active NF-κB transcription factor subunits to the nucleus and initiate the expression of its target genes (26, 27). To characterize the specific target of pD345L, several factors, including TRAF6, TAK1/TAB1, TBK1, IKKα and IKKβ that activate NF-κB at different steps of the signaling cascade, were tested. Dual luciferase reporter assay showed that pD345L was able to abolish NF-κB-Luc activity induced by TRAF6 (Figure 3A) and TAK1/TAB1 (Figure 3B) or significantly suppress NF-κB-Luc activity induced by TBK1 (Figure 3C), IKKα (Figure 3D) or IKKβ (Figure 3E). However, pD345L didn’t show inhibition on NF-κB-Luc activity induced by p65 overexpression (Figure 3F). Therefore, it is likely that pD345L modulates the NF-κB signaling at or downstream of IKK complex, while upstream of p65.

**FIG 3.**
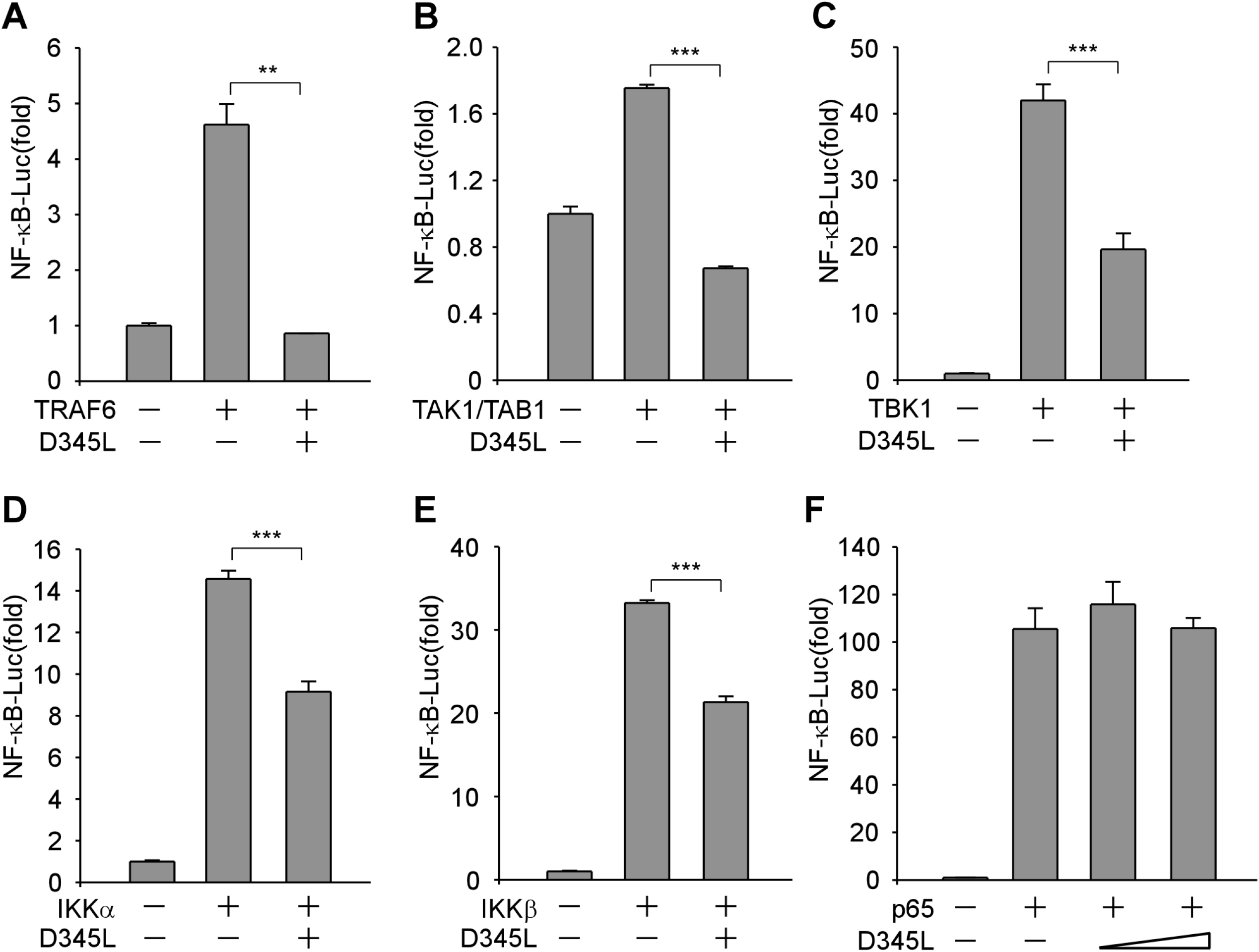
pD345L suppresses NF-κB activation at or downstream of IKKα/β and upstream of p65. 293T cells were transfected with NF-κB-Luc, pCMV-RL, and pcDNA3-Flag-TRAF6 (A), pcDNA4-HA-TAK1/TAB1 (B), pcDNA4-HA-TBK1 (C), pcDNA4-HA-IKKα (D), pcDNA4-HA-IKKβ (E), pcDNA4-HA-p65 (F), along with p3XFLAG-CMV-D345L or empty vector. The cells were collected 24 hours post transfection and luciferase activity was measured.

### pD345L interacts with IKKα and IKKβ

Activation of NF-κB signaling involves the phosphorylation of IκBα by IKK complex, which includes two catalytic subunits, IKKα and IKKβ, and one regulatory subunit IKKγ (28). According to the above results, we further explored whether pD345L directly targets IKK to inhibit IκBα phosphorylation. Co-immunoprecipitation assay was performed to analyze the protein-protein interaction between pD345L and IKK by co-transfection of Flag tagged D345L and HA tagged IKKα or IKKβ in 293T cells. Results showed the presence of HA-IKKα (Figure 4A) or HA-IKKβ (Figure 4C) in the Flag antibody immunoprecipitated pD345L protein complex and conversely the presence of Flag-pD345L in the HA antibody immunoprecipitated IKKα (Figure 4B) or IKKβ (Figure 4D) protein complex. IKKα and IKKβ subunits share 52% identity in their amino acid sequences and have similar primary structures that contain an N-terminal kinase domain (KD), a leucine zipper (LZ) domain and a C-terminal helix-loop-helix (HLH) domain (Figure 4E and 4F). To determine which domain of IKKα or IKKβ interacts with pD345L, several Flag tagged truncates containing KD, LZ or HLH domain of each subunit were constructed as previously described (29) (Figure 4E and F). Co-immunoprecipitation assay was performed and showed that HA tagged pD345L was detected in Flag antibody precipitated KD or HLH domain of IKKα (Figure 4G), and LZ domain of IKKβ (Figure 4H), which suggested that pD345L might target IKKα and IKKβ to suppress NF-κB signaling.

**FIG 4.**
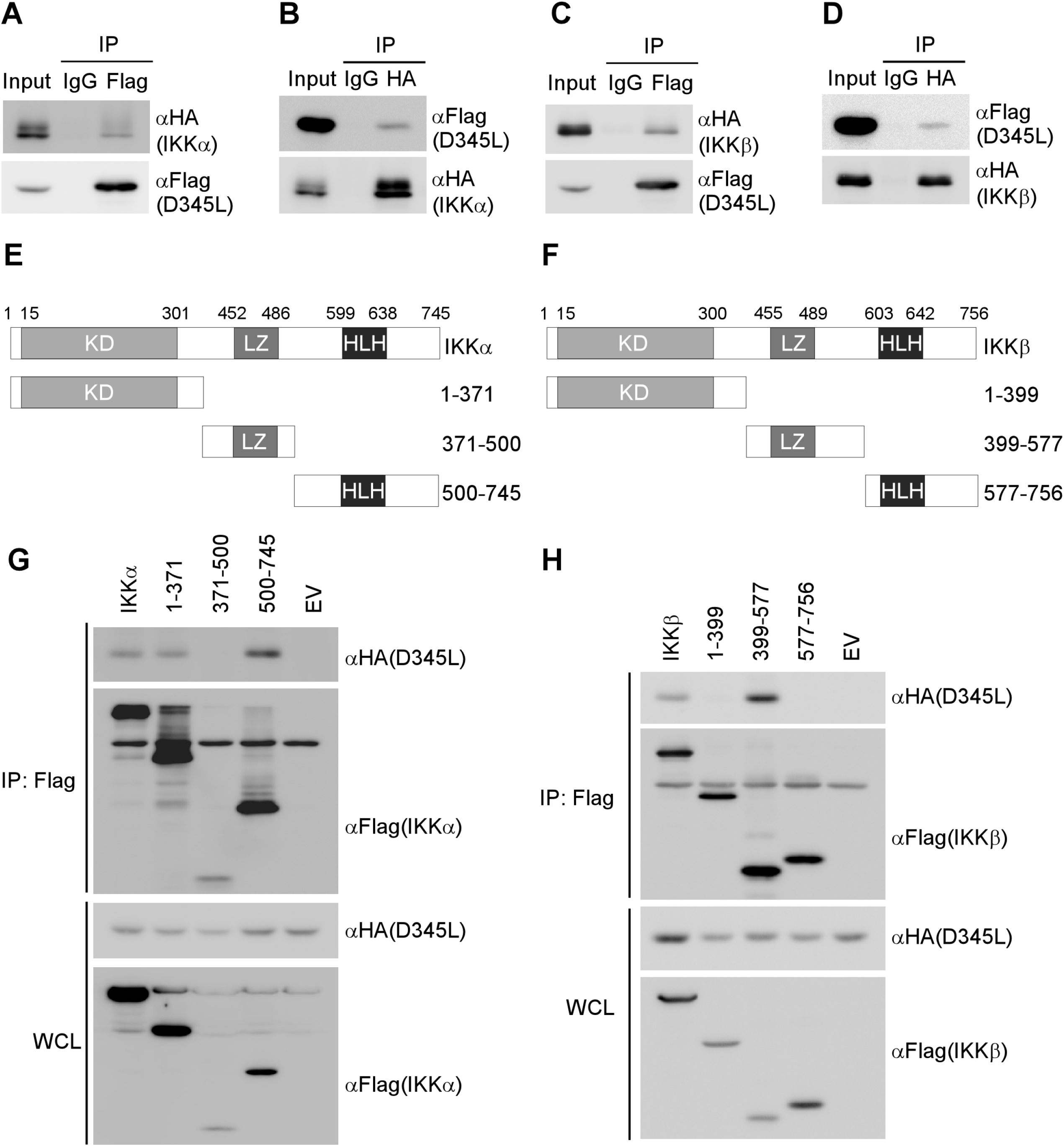
pD345L interacts with IKKα and IKKβ. Co-immunoprecipitaion assay was performed with whole cell lysates prepared with 293T cells co-transfected with Flag-D345L and HA-IKKα for 36 hours with Flag antibody (A), HA antibody (B) or control IgG. The immunocomplexs were analyzed by immunoblotting with the indicated antibodies. (C) and (D) The experiment was performed as for panel A or B except using HA-IKKβ. (E) and (F) Diagrams of the IKKα and IKKβ truncates. Both IKKα and IKKβ contain an N-terminal kinase domain (KD), a C-terminal helix– loop–helix domain (HLH) and a leucine zipper domain (LZ) as indicated. The experiment was performed as for panel A, except that HA-D345L and Flag-tagged IKKα, IKKα (1-371), IKKα (371-500), IKKα (500-745), empty vector (G), or Flag-tagged IKKβ, IKKβ (1-399), IKKβ (399-577), IKKβ (577-756), empty vector (H) were transfected as indicated.

### pD345L recruits IKKα and IKKβ to suppress their kinase activity on IκBα

It was showed that KD, LZ and HLH defective mutants of IKKα and IKKβ retain little or no IκBα activity, and either IKKα or IKKβ alone is able to mediate IκBα phosphorylation (30). Since pD345L interacts with KD and HLH domains of IKKα, LZ domain of IKKβ, we suspect that pD345L may directly target IKK to suppress their kinase activity on IκBα. To address this possibility, the kinase assay was performed with purified pD345L, IκBα, IKKα and IKKβ obtained by immunoprecipitation with or without ATP that provides a phosphate group necessary for the phosphorylation reaction to determine the newly formed p-IκBα. Results showed that in the absence of ATP, almost no p-IκBα was detected, while in the presence of ATP, p-IκBα was obviously increased when reacting with IKKα or IKKβ, but the increase of p-IκBα was prohibited by pD345L (Figure 5A). In addition, upon treatment with an increasing dose of pD345L, the IKKα or IKKβ activated p-IκBα was gradually decreased (Figure 5B and 5C). To further confirm these results, a D345L expression vector or empty vector was transfected into WSL cells followed by TNFα stimulation, immunoblotting results showed that p-IκBα was decreased at 0 and 30 minutes post TNFα treatment, and conversely the total IκBα level was increased at 0 and 60 minutes post TNFα treatment (Figure 5D). This suggests that pD345L overexpression inhibited TNFα-induced IκBα phosphorylation and thus the total IκBα level was recovered, which would enhance the interaction between IκBα and p65. Therefore, p65 translocation was examined in WSL cells with or without pD345L expression. Results showed that p65 translocated from cytoplasm to the nucleus after TNFα treatment, however it remained in the cytoplasm of cells expressing pD345L (Figure 5E). Therefore, these findings indicate that pD345L directly inhibits IκBα phosphorylation through association with IKKα and IKKβ, further suppressing p65 nuclear translocation.

**FIG 5.**
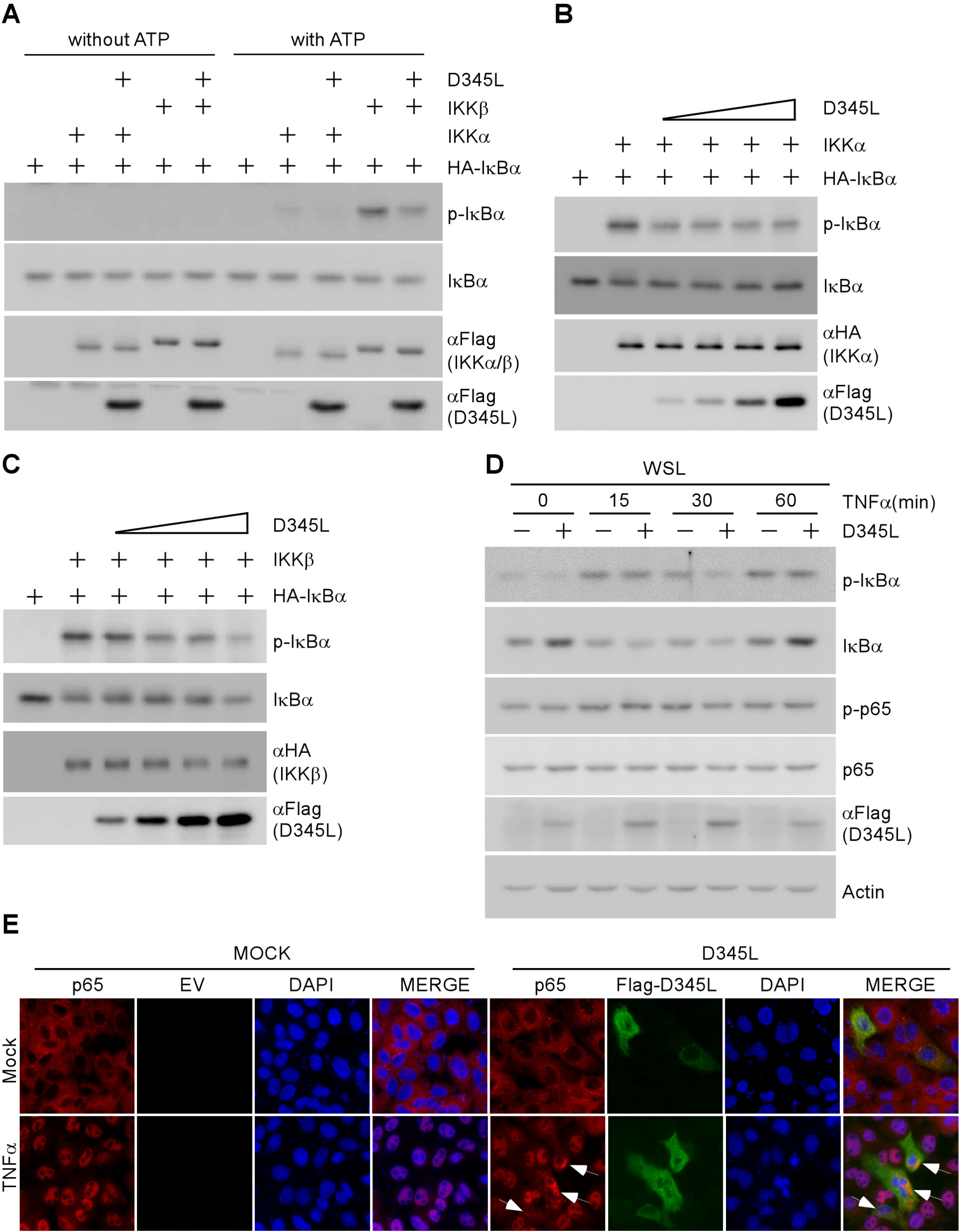
pD345L targets IKKα/β to inhibit their kinase activity on IκBα. (A) 293T cells were transfected with HA-IκBα, Flag-IKKα, Flag-IKKβ or Flag-D345L respectively, 36 hours post transfection, these proteins were immunoprecipitated with the anti-Flag or anti-HA antibody respectively. Kinase assay of the immunoprecipitated proteins in vitro was carried out with or without ATP, and analyzed by immunoblotting. (B) and (C) The experiment was performed as for panel A except using an increasing dose of pD345L in the reaction. (D) WSL cells were transfected with Flag-D345L or empty vector for 30 hours followed by treatment with TNFα (40ng/ml) at the indicated time points. Cells were collected for immunoblotting with the indicated antibodies. (E) WSL cells were transfected with Flag-D345L or empty vector for 36 hours followed by treatment with TNFα (40ng/ml) for 30 minutes and stained with anti-Flag (green) or anti-p65 (red) antibody. The nuclei were stained with DAPI (blue) and images were acquired with a Nikon fluorescence microscope.

### pD345L inhibits the NF-κB signaling independent on its exonuclease activity

pD345L protein contains an N-terminal 5’ → 3’ exonuclease domain and shows strong similarity with lambda phage exonuclease, while its C-terminal function remains unknown (12). To investigate whether inhibition of NF-κB activity relies on pD345L exonuclease activity, D345L catalytic mutant (D345L-CM) was constructed (Figure 6A). D345L-CM was replaced Aspartic acid 108 (D108), Glutamic acid 144 (E144), and lysine 146 (K146) residues into Alanine (A) according to lambda phage exonuclease catalytic center residues (31). A dual-luciferase reporter assay was conducted with transfection of D345L or D345L-CM along with cGAS/STING or TBK1, and showed that pD345L-CM was still able to suppress the activation of IFNβ promoter (Figure 6B and 6C) and NF-κB-Luc (Figure 6D). Consistent with these, stimulation of IFNβ mRNA expression by 2’3’cGAMP was attenuated by both pD345L and pD345L-CM in WSL cells (Figure 6E). To confirm these results, D345L N- or C-terminal truncates with or without the exonuclease domain (D345L-N and D345L-C) were constructed. Unfortunately, D345L-N couldn’t be expressed in 293T cells. Therefore, D345L-C was used for NF-κB luciferase reporter assay and RT-PCR. It showed that pD345L-C still capable of inhibiting on the NF-κB signaling (Figure 6F) and IFNβ production (Figure 6G). Finally, co-immunoprecipitation assay was performed to detect pD345L-C and IKK interaction. Results showed that HA-IKKα and HA-IKKβ was detected in Flag antibody precipitated pD345L or pD345L-C (Figure 6H and 6I).

**FIG 6.**
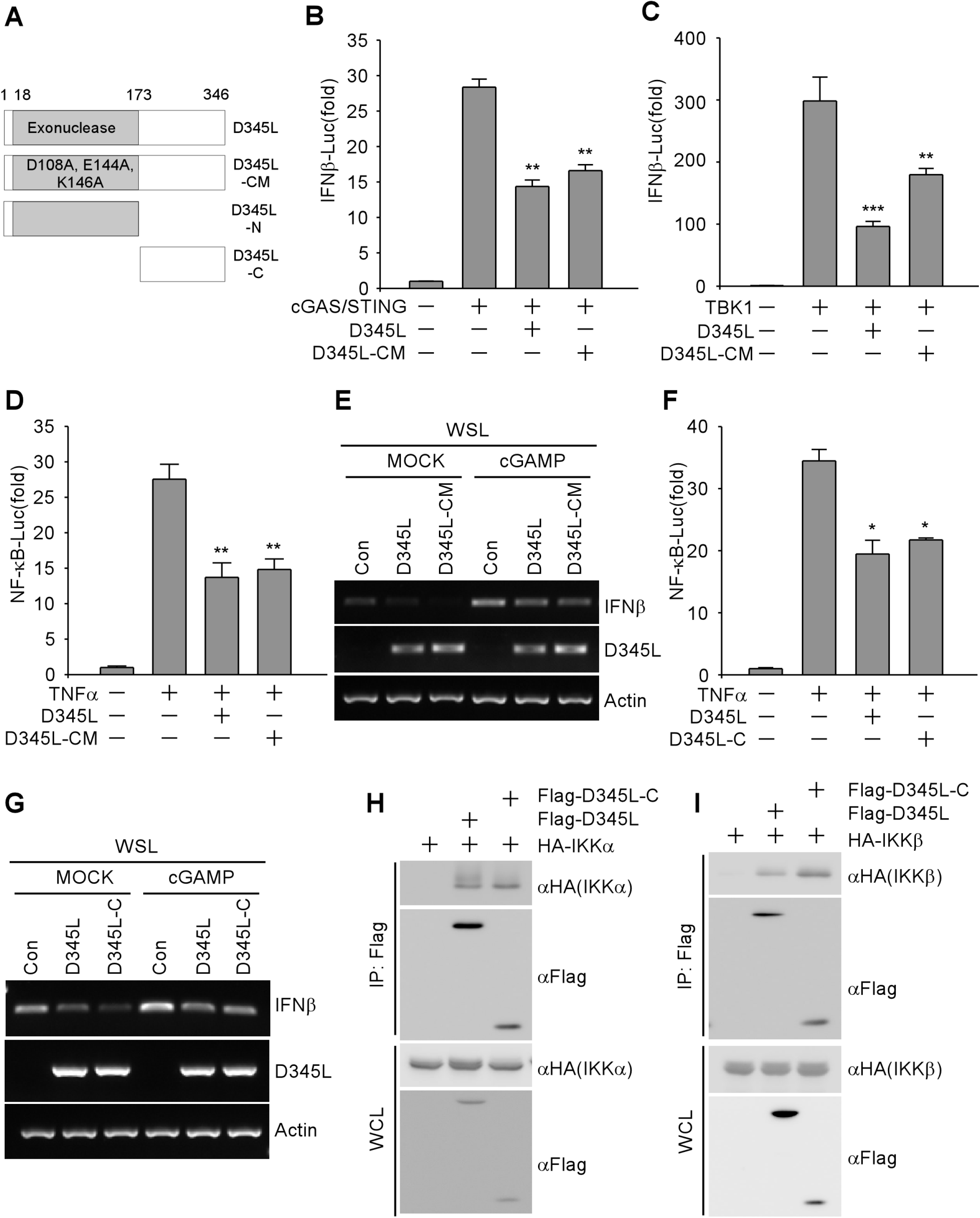
pD345L inhibition of NF-κB signaling was independent of its exonuclease activity. (A) A schematic representation of the D345L catalytic activity mutant (D345L-CM), N-terminal mutant (D345L-N), and N-terminal deletion mutant (D345L-C). (B) 293T cells transfected with IFNβ-Luc, pCMV-RL, pcDNA3.1-HA-cGAS and pcDNA3-2xFlag-STING, along with p3XFLAG-CMV-D345L, p3XFLAG-CMV-D345L-CM or empty vector for 24 hours, were collected for measuring the luciferase activity. (C) The experiment was performed as in panel B except using pcDNA4-HA-TBK1. (D) 293T cells transfected with NF-κB-Luc and pCMV-RL along with p3XFLAG-CMV-D345L, p3XFLAG-CMV-D345L-CM or empty vector for 24 hours followed by treatment with TNFα (40ng/ml) for 6 hours, were collected for measuring the luciferase activity. (E) WSL cells transfected with p3XFLAG-CMV-D345L or p3XFLAG-CMV-D345L-CM or empty vector for 30 hours followed by stimulation with 2’3’ cGAMP (5μg/ml) for 4 hours and harvested for RNA extraction. Semiquantitative RT-PCR assay was carried out to detect IFNβ, D345L and Actin mRNA expression levels. (F) The experiment was performed as for panel D, except that p3XFLAG-CMV-D345L-C was transfected. (G) The experiment was performed as for panel E, except that p3XFLAG-CMV-D345L-C was transfected. 293T cells were co-transfected with Flag-D345L, Flag-D345L -C and HA-IKKα (H), HA-IKKβ (I) or empty vector for 36 hours, and the whole cell lysates were immunoprecipitated with Flag antibody. The immunocomplexs were analyzed by immunoblotting with the indicated antibodies.

Taken together, ASFV pD345L protein blocks NF-κB signaling via interacting with IKK and subsequently suppressing their kinase activity on IκBα.

## Discussion

Since expanding of ASFV affected areas into Asian countries, especially China, an effective vaccine is urgently needed more than ever. However, development of an effective vaccine is largely hindered due to limited knowledge of protective antigens and the interaction between ASFV infection and host immune responses. The NF-κB signaling plays a critical role in innate immune system by controlling transcription of antiviral genes, including IFNs, cytokines, chemokines as well as their modulators and immunoreceptors, and genes involved in cell cycle, apoptosis and stress response. Therefore, viruses have evolved multiple strategies to disrupt the NF-κB signaling cascade in order to guarantee efficient replication. Kinase activity of IKK protein complex (IKKα/β/γ) is essential for NF-κB activation and thus becomes a common target of many viruses. For examples, Influenza A virus NS1 protein targets kinase domain (KD) of IKKα/β, blocking IKKβ-mediated phosphorylation and degradation of IκBα in canonical pathway and IKKα-mediated processing of precursor protein p100 to p52 in non-canonical pathway (32). In addition, Vaccinia Virus (VACV) B14 protein inhibits IKKβ activation through binding to its KD and scaffold and dimerization domain (SDD) (33-35). Moreover, Molluscum contagiosum virus (MCV) MC160 protein is capable of inhibiting IKK complex formation (36).

In this study, we found that ASFV-encoded lambda-like exonuclease, pD345L, is an inhibitor of cGAS/STING-NF-κB signaling. As a result, overexpression of pD345L obviously suppressed cGAMP-induced transcript levels of IFNβ and inflammatory cytokines (IL-1α, IL-6, IL-8, and TNFα) (Figure 2). Importantly, we demonstrated that along the NF-κB signaling cascade, pD345L specifically targets the IKK complex through interacting with KD and HLH domains of IKKα (Figure 4G), and LZ domain of IKKβ (Figure 4H). Previous reports showed that LZ domain is required for proper formation of IKKα and IKKβ homo- or hetero-dimers, and LZ domain or HLH motif defective mutants of IKKα and IKKβ block their kinase activities (30, 37, 38). Interestingly, we found that in the presence of pD345L, IκBα phosphorylation by IKK was decreased (Figure 5A). To our knowledge, this is the first time to show that an ASFV protein is able to block IKKα/β kinase activity through protein-protein interaction.

In addition to NF-κB-dependent targets, IKK also regulates a number of substrates involved in regulating cell growth, apoptosis, autophagy and metabolism (39). For examples, on the one hand, IKKβ promotes transcription of anti-apoptotic genes in NF-κB canonical pathway; on the other hand, IKK inhibits apoptosis by phosphorylating pro-apoptotic protein PUMA at serine10 and Bad at serine26 (40, 41). In addition, active IKK subunits stimulate autophagy and knockout of IKKβ inhibits the activation of autophagy in mice (42). Autophagy related proteins, ATG16L1 (autophagy-related 16-like 1) and AMBRA1 (autophagy and beclin-1 regulator 1) were identified to be IKK substrates (43, 44). It has been shown that precise manipulation of host cell apoptosis by ASFV is critical to accomplish different infection stages of its life cycle (45-47). However, autophagy is not required for ASFV replication and autophagosome formation is inhibited upon ASFV infection (48). Therefore, through modulating IKK activity, pD345L possibly regulates multiple biological pathways. Whether pD345L suppresses IKK kinase activity on PUMA, Bad or other substrates to regulate apoptosis or autophagy in ASFV infected cells will be worthwhile to be further explored.

To determine the effect of D345L deletion on virus-induced IFNβ response, generation of D345L-deficient ASFV was tried but failed, consistent with a previous report (12), suggesting the essential role of pD345L in ASFV replication. Conditional deletion of D345L will be tried in further studies(49, 50). pD345L is a lambda-like exonuclease and such proteins also found in phycodnaviruses, mimivirus and bacteriophages are implicated in DNA repair and chromosome recombination (4, 31). Conventional vaccine development approaches are proven to be inapplicable to ASFV (51, 52). Neither subunit or DNA vaccine provides efficient protection (53). Gene deleted live-attenuated vaccine (LAV) candidates render adequate protection by eliciting both humoral and cellular immunity, but establishment of chronic or persistent infection in vaccinated animals and risk of recombination with filed strains are unsolved key issues. Here, we showed that D345L is not only an essential gene for ASFV replication but also inhibits host immune response, which may explain why LAVs will maintain the immune evasion ability but to a lesser extent. To overcome LAV-induced persistent infection, one potential strategy would be generation of replication-defective viruses, limiting or abrogating viral replication in animals by targeting replication essential viral genes. Given the dual role of pD345L in virus replication and immune evasion, it may serve as a potential target for replication-defective virus vaccine development.

## Materials and Methods

### Plasmids

For mammalian expression, p3XFLAG-CMV-D345L was kindly provided by Dr Hongjun Chen (Shanghai Veterinary Research Institute). The pcDNA3.1-HA-cGAS and pcDNA3-2xFlag-STING were generated as previous described(54). Pig IKKα (EU399820.1), IKKβ (NM_001099935), TAB1 (NM_001244067), TBK1(NM_001105292), IκBα(NM_001005150), TAK1 (KU504629.1), p65 (KC316023.1) were amplified from cDNA of PK15 and cloned into pcDNA4-HA. D345L catalytic mutant (D345L-D108A, E144A and K146A) was constructed using homologous recombination kit (Vazyme Biotech, China). The luciferase reporter plasmids used in this study were pGL3-Basic-IFN-β-Luc, pGL4.32-NF-κB-Luc and pGluc-IRF3-Luc, which contain the -296 to +52 fragment of the pig IFN-β promoter (54), five copies of NF-κB-response element and four copies of IRF3-response element, respectively.

### Cells, Antibodies and Reagents

293T and PK15 cells were grown in Dulbecco’s modified Eagle medium (DMEM; Gibco-BRL) supplemented with 8% fetal bovine serum (PAN-Biotech, Dorset, UK) and 1% of penicillin and streptomycin (Beyotime Biotechnology, Shanghai, China) at 37 ° C in a 5% CO_2_ incubator. WSL cells were maintained in RPMI 1640 Medium (Gibco-BRL) supplemented with 10% fetal bovine serum and 1% of penicillin and streptomycin.

The primary antibodies used in this study from Cell Signaling (Beverly, MA) include rabbit anti-phospho-p65 (serine 536), rabbit anti-p65, mouse anti-IκBα, rabbit anti-p-IκBα (Serine 32), rabbit anti-IKKα, rabbit anti-IKKβ, rabbit anti-hemagglutinin (HA) antibodies. Mouse anti-Flag antibody was from Sigma-Aldrich (St. Louis, MO, USA) and rabbit anti-actin antibody was from Proteintech (Wuhan, China). HRP-conjugated goat anti-mouse IgG (H + L) or anti-rabbit IgG (H + L) were purchased from Millipore. Alexa 488-conjugated goat anti-mouse IgG antibody and Alexa 555-conjugated goat anti-rabbit IgG antibody were purchased from Thermo Fisher Scientific (MA, USA). 2’3’ cGAMP and poly(I:C) HMW were purchased from Invivogen (San Diego, CA, USA). Recombinant TNFα was purchased from Sigma-Aldrich (St. Louis, MO, USA). The protease inhibitor cocktail was purchased from Thermo Fisher Scientific (MA, USA).

### RNA isolation and Semi-Quantitative PCR

Total RNA was extracted from 293T, WSL and PK15 cells using Simple P Total RNA Extraction Kit (Bioer Technology, China) according to the manufacturer’s instructions. Samples were subjected to reverse transcription using HiScript II Q RT kit (Vazyme Biotech, China). The cDNA was used as a template for Semi-quantitative RT-PCR to investigate the mRNA levels of pig IL-1α, IL-6, IL-8, IFNβ, TNFα, D345L and Actin. The mRNA levels of IFNβ and GAPDH were quantitated by SYBR green-based quantitative real-time PCR (Vazyme Biotech, China) using a Life Technology instrument. The primers used here were listed in Table 1.

**Table 1.**
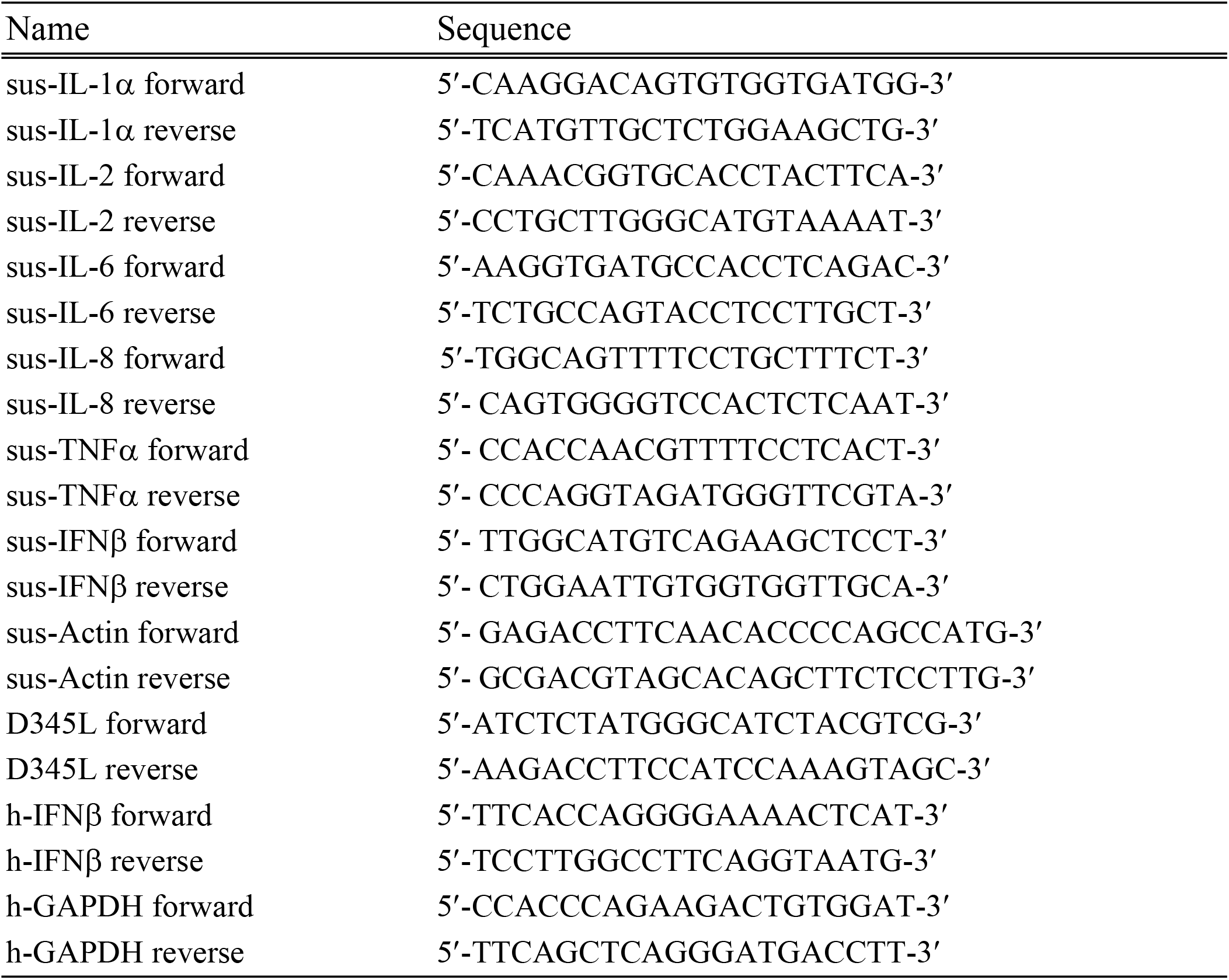
The primers used for RT-PCR and Q-RT-PCR.

### Transfection and Dual-Luciferase assay

The dual-luciferase assay was performed in triplicate according to the manufacturer’s instructions. 293T cells were seeded on 24-well plates (Thermo Scientific) at a density of 1×10^5^ cells per well overnight before transfection. Cells were then co-transfected with p3XFLAG-CMV-D345L (200ng), pGL3-Basic-IFN-β-Luc (200ng) and an internal control Renilla luciferase vector pCMV-RL (2ng) for 24 hours using Lipofectamine 2000 reagent (Thermo Fisher Scientific, MA, USA). Luciferase activity was measured with a dual-luciferase assay kit (Promega, Madison, WI, USA) and a MD SpectraMax iD5 instrument.

### Co-Immunoprecipitation and Immunoblotting

293T cells were co-transfected with p3XFLAG-CMV-D345L (1.5μg) and pcDNA4-HA-IKKα/β (1.5μg). After 36 hours transfection, cells were washed with ice-cold PBS and lysed with lysis buffer (50 mM Tris, pH 7.5, 150 mM NaCl, 1 mM EDTA, and 1% NP-40 supplemented with protease inhibitor) and incubated with anti-Flag or anti-HA antibody, together with protein A/G magnetic beads for 2 hours at 4 ° C. After 8 times washes with ice-cold lysis buffer, proteins were eluted with SDS sample buffer and analyzed by immunoblotting. For immunoblotting, the co-immunoprecipitation sample and 2% whole-cell lysates were analyzed by SDS-PAGE and transferred to a polyvinylidene difluoride (PVDF) membrane (Pall Corp.). The membrane was blocked in 3% skim milk in phosphate buffered saline with Tween 20 (PBST) for 1 hour and then incubated with primary antibodies at 4 °C overnight and anti-mouse or anti-rabbit IgG antibody conjugated to HRP.

### Kinase assay

293T cells were transfected with plasmids expressing the Flag-tagged or HA-tagged proteins. 36 hours post-transfection, the cell lysates immunoprecipitated with the anti-flag or anti-HA antibody were subjected to the kinase assay in a reaction buffer (25 mM Tris-HCl, pH 7.5, 0.01% TritonX-100, 10 mM MgCl_2_, 0.5 mM Na_3_VO_4_, 2.5 mM DTT, 0.5 mM EGTA, and 50 μM ATP) with or without ATP at 30°C for 1 hour. The kinase reaction products were analyzed by immunoblotting.

### Immunofluorescence assay

WSL cells were grown on coverslips and transfected with p3XFLAG-CMV-D345L or empty vector for 36 hours followed by treatment with TNFα (40ng/ml) for 30 minutes. The cells were fixed and permeabilized with 4% formaldehyde and 0.1% Triton X-100 at 37 °C for 30 minutes. After washing with glycine-PBS, cells were blocked with 3% BSA in PBS for 1 hour at room temperature. The slides were incubated with primary antibody (1:200) for 1 hour, followed by secondary antibody (1:500) for 30 minutes. The nuclei were stained with DAPI and images were obtained with Nikon fluorescence microscope (TS100-F; DSRi2).

### Statistical analysis

All experiments were performed at least three times unless otherwise indicated. Data are presented as the means ± standard deviations (SDs). Statistical significance between groups was determined using Student’s t-test in GraphPad Prism 7.0 software (La Jolla, CA, USA). *P < 0.05, **P < 0.01, ***P < 0.001.

## Data availability

All of the data are fully available without restriction.

## Acknowledgments

This work was supported by the National Key Research and Development Program of China (2017YFD0502306-1), the National Natural Science Foundation of China (U19A2039 and 31900141), Key Research and Development Program of Jiangsu Province (BE2020407), the Fundamental Research Funds for the Central Universities (KJQN202022), and the Priority Academic Program Development of Jiangsu Higher Education Institutions (PAPD).

## Author contributions

Huan Chen, Yong-Sam Jung, Xiaodong Wu, and Yingjuan Qian designed the study and analyzed data. Huan Chen and Zhenzhong Wang performed the experiments with the help of Xiaoyu Gao, Jiaxuan Lv and Yongxin Hu. Yingjuan Qian, Yong-Sam Jung, Xiaodong Wu, Shanyuan Zhu, Jianjun Dai developed the concept and interpreted data. Huan Chen, Yingjuan Qian and Yong-Sam Jung drafted the manuscript, and all authors commented on it. Yingjuan Qian supervised the entire project.

## References

1. Galindo I, Alonso C. 2017. African Swine Fever Virus: A Review. Viruses 9.

2. Mighell E, Ward MP. 2021. African Swine Fever spread across Asia, 2018-2019. Transbound Emerg Dis doi:10.1111/tbed.14039.

3. Vilanova E, Tovar AMF, Mourao PAS. 2019. Imminent risk of a global shortage of heparin caused by the African Swine Fever afflicting the Chinese pig herd. J Thromb Haemost 17:254–256.

4. Iyer LM, Aravind L, Koonin EV. 2001. Common origin of four diverse families of large eukaryotic DNA viruses. J Virol 75:11720–34.

5. Iyer LM, Balaji S, Koonin EV, Aravind L. 2006. Evolutionary genomics of nucleo-cytoplasmic large DNA viruses. Virus Res 117:156–84.

6. Dixon LK, Chapman DA, Netherton CL, Upton C. 2013. African swine fever virus replication and genomics. Virus Res 173:3–14.

7. de Villiers EP, Gallardo C, Arias M, da Silva M, Upton C, Martin R, Bishop RP. 2010. Phylogenomic analysis of 11 complete African swine fever virus genome sequences. Virology 400:128–36.

8. Suzuki S, Kameoka M, Nakaya T, Kimura T, Nishi N, Hirai K, Ikuta K. 1997. Superoxide generation by monocytes following infection with human cytomegalovirus. Immunopharmacology 37:185–90.

9. Li Z, Xu X, Leng X, He M, Wang J, Cheng S, Wu H. 2017. Roles of reactive oxygen species in cell signaling pathways and immune responses to viral infections. Arch Virol 162:603–610.

10. Cobbold C, Windsor M, Parsley J, Baldwin B, Wileman T. 2007. Reduced redox potential of the cytosol is important for African swine fever virus capsid assembly and maturation. J Gen Virol 88:77–85.

11. Redrejo-Rodriguez M, Garcia-Escudero R, Yanez-Munoz RJ, Salas ML, Salas J. 2006. African swine fever virus protein pE296R is a DNA repair apurinic/apyrimidinic endonuclease required for virus growth in swine macrophages. J Virol 80:4847–57.

12. Modesto Redrejo-Rodríguez JMRg, José Salas and María L. Salas 2011. DNA Repair - On the Pathways to Fixing DNA Damage and Errors., Repair of Viral Genomes by Base Excision Pathways: African Swine Fever Virus as a Paradigm doi:10.5772/871.InTech.

13. Bonizzi G, Karin M. 2004. The two NF-kappa B activation pathways and their role in innate and adaptive immunity. Trends in Immunology 25:280–288.

14. Anest V, Hanson JL, Cogswell PC, Steinbrecher KA, Strahl BD, Baldwin AS. 2003. A nucleosomal function for IkappaB kinase-alpha in NF-kappaB-dependent gene expression. Nature 423:659–63.

15. Yamamoto Y, Verma UN, Prajapati S, Kwak YT, Gaynor RB. 2003. Histone H3 phosphorylation by IKK-alpha is critical for cytokine-induced gene expression. Nature 423:655–9.

16. Xiao G, Harhaj EW, Sun SC. 2001. NF-kappaB-inducing kinase regulates the processing of NF-kappaB2 p100. Mol Cell 7:401–9.

17. Revilla Y, Callejo M, Rodriguez JM, Culebras E, Nogal ML, Salas ML, Vinuela E, Fresno M. 1998. Inhibition of nuclear factor kappaB activation by a virus-encoded IkappaB-like protein. J Biol Chem 273:5405–11.

18. Granja AG, Perkins ND, Revilla Y. 2008. A238L inhibits NF-ATc2, NF-kappa B, and c-Jun activation through a novel mechanism involving protein kinase C-theta-mediated up-regulation of the amino-terminal transactivation domain of p300. J Immunol 180:2429–42.

19. Correia S, Ventura S, Parkhouse RM. 2013. Identification and utility of innate immune system evasion mechanisms of ASFV. Virus Res 173:87–100.

20. Wang X, Wu J, Wu Y, Chen H, Zhang S, Li J, Xin T, Jia H, Hou S, Jiang Y, Zhu H, Guo X. 2018. Inhibition of cGAS-STING-TBK1 signaling pathway by DP96R of ASFV China 2018/1. Biochem Biophys Res Commun 506:437–443.

21. Li D, Yang W, Li L, Li P, Ma Z, Zhang J, Qi X, Ren J, Ru Y, Niu Q, Liu Z, Liu X, Zheng H. 2021. African Swine Fever Virus MGF-505-7R Negatively Regulates cGAS-STING-Mediated Signaling Pathway. J Immunol 206:1844–1857.

22. Reis AL, Abrams CC, Goatley LC, Netherton C, Chapman DG, Sanchez-Cordon P, Dixon LK. 2016. Deletion of African swine fever virus interferon inhibitors from the genome of a virulent isolate reduces virulence in domestic pigs and induces a protective response. Vaccine 34:4698–4705.

23. Afonso CL, Piccone ME, Zaffuto KM, Neilan J, Kutish GF, Lu Z, Balinsky CA, Gibb TR, Bean TJ, Zsak L, Rock DL. 2004. African swine fever virus multigene family 360 and 530 genes affect host interferon response. J Virol 78:1858–64.

24. Garcia-Belmonte R, Perez-Nunez D, Pittau M, Richt JA, Revilla Y. 2019. African Swine Fever Virus Armenia/07 Virulent Strain Controls Interferon Beta Production through the cGAS-STING Pathway. J Virol 93.

25. Lawrence T. 2009. The nuclear factor NF-kappaB pathway in inflammation. Cold Spring Harb Perspect Biol 1:a001651.

26. Balka KR, Louis C, Saunders TL, Smith AM, Calleja DJ, D’Silva DB, Moghaddas F, Tailler M, Lawlor KE, Zhan Y, Burns CJ, Wicks IP, Miner JJ, Kile BT, Masters SL, De Nardo D. 2020. TBK1 and IKKepsilon Act Redundantly to Mediate STING-Induced NF-kappaB Responses in Myeloid Cells. Cell Rep 31:107492.

27. Liu T, Zhang L, Joo D, Sun SC. 2017. NF-kappaB signaling in inflammation. Signal Transduct Target Ther 2.

28. Karin M. 1999. How NF-kappaB is activated: the role of the IkappaB kinase (IKK) complex. Oncogene 18:6867–74.

29. Cui J, Zhu L, Xia X, Wang HY, Legras X, Hong J, Ji J, Shen P, Zheng S, Chen ZJ, Wang RF. 2010. NLRC5 negatively regulates the NF-kappaB and type I interferon signaling pathways. Cell 141:483–96.

30. Zandi E, Chen Y, Karin M. 1998. Direct phosphorylation of IkappaB by IKKalpha and IKKbeta: discrimination between free and NF-kappaB-bound substrate. Science 281:1360–3.

31. Kovall R, Matthews BW. 1997. Toroidal structure of lambda-exonuclease. Science 277:1824–7.

32. Gao S, Song L, Li J, Zhang Z, Peng H, Jiang W, Wang Q, Kang T, Chen S, Huang W. 2012. Influenza A virus-encoded NS1 virulence factor protein inhibits innate immune response by targeting IKK. Cell Microbiol 14:1849–66.

33. Benfield CT, Mansur DS, McCoy LE, Ferguson BJ, Bahar MW, Oldring AP, Grimes JM, Stuart DI, Graham SC, Smith GL. 2011. Mapping the IkappaB kinase beta (IKKbeta)-binding interface of the B14 protein, a vaccinia virus inhibitor of IKKbeta-mediated activation of nuclear factor kappaB. J Biol Chem 286:20727–35.

34. Tang Q, Chakraborty S, Xu G. 2018. Mechanism of vaccinia viral protein B14-mediated inhibition of IkappaB kinase beta activation. J Biol Chem 293:10344–10352.

35. Chen RA, Ryzhakov G, Cooray S, Randow F, Smith GL. 2008. Inhibition of IkappaB kinase by vaccinia virus virulence factor B14. PLoS Pathog 4:e22.

36. Nichols DB, Shisler JL. 2006. The MC160 protein expressed by the dermatotropic poxvirus molluscum contagiosum virus prevents tumor necrosis factor alpha-induced NF-kappaB activation via inhibition of I kappa kinase complex formation. J Virol 80:578–86.

37. Zandi E, Rothwarf DM, Delhase M, Hayakawa M, Karin M. 1997. The IkappaB kinase complex (IKK) contains two kinase subunits, IKKalpha and IKKbeta, necessary for IkappaB phosphorylation and NF-kappaB activation. Cell 91:243–52.

38. Kwak YT, Guo J, Shen J, Gaynor RB. 2000. Analysis of domains in the IKKalpha and IKKbeta proteins that regulate their kinase activity. J Biol Chem 275:14752–9.

39. Antonia RJ, Hagan RS, Baldwin AS. 2021. Expanding the View of IKK: New Substrates and New Biology. Trends Cell Biol 31:166–178.

40. Sandow JJ, Jabbour AM, Condina MR, Daunt CP, Stomski FC, Green BD, Riffkin CD, Hoffmann P, Guthridge MA, Silke J, Lopez AF, Ekert PG. 2012. Cytokine receptor signaling activates an IKK-dependent phosphorylation of PUMA to prevent cell death. Cell Death Differ 19:633–41.

41. Yan J, Xiang J, Lin Y, Ma J, Zhang J, Zhang H, Sun J, Danial NN, Liu J, Lin A. 2013. Inactivation of BAD by IKK inhibits TNFalpha-induced apoptosis independently of NF-kappaB activation. Cell 152:304–15.

42. Criollo A, Senovilla L, Authier H, Maiuri MC, Morselli E, Vitale I, Kepp O, Tasdemir E, Galluzzi L, Shen S, Tailler M, Delahaye N, Tesniere A, De Stefano D, Younes AB, Harper F, Pierron G, Lavandero S, Zitvogel L, Israel A, Baud V, Kroemer G. 2010. The IKK complex contributes to the induction of autophagy. EMBO J 29:619–31.

43. Di Rita A, Peschiaroli A, P Da, Strobbe D, Hu Z, Gruber J, Nygaard M, Lambrughi M, Melino G, Papaleo E, Dengjel J, El Alaoui S, Campanella M, Dotsch V, Rogov VV, Strappazzon F, Cecconi F. 2018. HUWE1 E3 ligase promotes PINK1/PARKIN-independent mitophagy by regulating AMBRA1 activation via IKKalpha. Nat Commun 9:3755.

44. Diamanti MA, Gupta J, Bennecke M, De Oliveira T, Ramakrishnan M, Braczynski AK, Richter B, Beli P, Hu Y, Saleh M, Mittelbronn M, Dikic I, Greten FR. 2017. IKKalpha controls ATG16L1 degradation to prevent ER stress during inflammation. J Exp Med 214:423–437.

45. Alonso C, Galindo I, Cuesta-Geijo MA, Cabezas M, Hernaez B, Munoz-Moreno R. 2013. African swine fever virus-cell interactions: from virus entry to cell survival. Virus Res 173:42–57.

46. Carrascosa AL, Bustos MJ, Nogal ML, Gonzalez de Buitrago G, Revilla Y. 2002. Apoptosis induced in an early step of African swine fever virus entry into vero cells does not require virus replication. Virology 294:372–82.

47. Hernaez B, Escribano JM, Alonso C. 2006. Visualization of the African swine fever virus infection in living cells by incorporation into the virus particle of green fluorescent protein-p54 membrane protein chimera. Virology 350:1–14.

48. Hernaez B, Cabezas M, Munoz-Moreno R, Galindo I, Cuesta-Geijo MA, Alonso C. 2013. A179L, a new viral Bcl2 homolog targeting Beclin 1 autophagy related protein. Curr Mol Med 13:305–16.

49. Garcia-Escudero R, Andres G, Almazan F, Vinuela E. 1998. Inducible gene expression from African swine fever virus recombinants: analysis of the major capsid protein p72. J Virol 72:3185–95.

50. Matamoros T, Alejo A, Rodriguez JM, Hernaez B, Guerra M, Fraile-Ramos A, Andres G. 2020. African Swine Fever Virus Protein pE199L Mediates Virus Entry by Enabling Membrane Fusion and Core Penetration. mBio 11.

51. Stone SS, Hess WR. 1967. Antibody response to inactivated preparations of African swine fever virus in pigs. Am J Vet Res 28:475–81.

52. Blome S, Gabriel C, Beer M. 2014. Modern adjuvants do not enhance the efficacy of an inactivated African swine fever virus vaccine preparation. Vaccine 32:3879–82.

53. Revilla Y, Perez-Nunez D, Richt JA. 2018. African Swine Fever Virus Biology and Vaccine Approaches. Adv Virus Res 100:41–74.

54. Bo Z, Miao Y, Xi R, Zhong Q, Bao C, Chen H, Sun L, Qian Y, Jung YS, Dai J. 2020. PRV UL13 inhibits cGAS-STING-mediated IFN-beta production by phosphorylating IRF3. Vet Res 51:118.

